# The Effect of Cognitive Load and Default Nudges on Decision-Making: Experimental Evidence from a Behavioural Study

**DOI:** 10.1101/2025.05.05.652071

**Authors:** Mercede Erfanian, Luc Meunier, Jean-Francois Gajewski

## Abstract

Cognitive overload can impair professional scepticism in high-stakes contexts such as auditing. In these settings, sustaining professional scepticism is essential. Default nudges, or pre-selected options, may offset these effects by reducing cognitive demands. We conducted two online experiments to examine how cognitive load and default nudges influence professional scepticism in auditing decisions. Experiment 1 validated a dot memory task manipulation of cognitive load and identified low and high load conditions for subsequent testing. Experiment 2 embedded this manipulation in Phillips’ audit task, used for measuring professional scepticism in audit. Results showed that cognitive load slowed responses and reduced accuracy. Default nudges accelerated responding and improved accuracy under load, but only when aligned with the most probable response; misaligned nudges reduced accuracy. These findings suggest that defaults act as conditional scaffolds under cognitive strain, supporting judgment and decision-making in some contexts but introducing risks in others. Misaligned defaults reduced accuracy, indicating that they can exploit intuitive responding rather than enhance deliberation.

## 1. Introduction

The brain, though only ∼2% of body weight, consumes ∼20% of resting energy (Raichle et al., 2002). This reflects the metabolic demands of executive functions, yet adaptive mechanisms balance energy use with sustained performance (Li et al., 2020; Padamsey et al., 2023; Raichle et al., 2002; Sterling et al., 2015). Efficiency is especially relevant during cognitively demanding tasks (Baumeister, 2001; Gailliot, 2008), such as decision-making (Wiehler et al., 2022), which relies on working memory (WM), cognitive control, and flexibility (Botvinick et al., 2004; Mansouri et al., 2017), all limited and energy-intensive capacities.

Scepticism is the disposition to question assumptions and critically evaluate evidence, supporting high-quality decisions in uncertain contexts. It enables inhibition of irrelevant or misleading inputs (Stanovich et al., 2016) and underpins *epistemic vigilance*, the evolved capacity to assess source reliability (Heyes, 2018; Mercier et al., 2017; Sperber et al., 2010). Without it, uncritical acceptance of claims undermines epistemic integrity and increases susceptibility to misinformation (Sperber et al., 2010). Thus, scepticism safeguards judgment and decision-making accuracy, especially where truth is consequential, and is central to reasoning under uncertainty with incomplete or ambiguous information (Bago et al., 2020; Gigerenzer et al., 2011; Kahneman et al., 2021; Lewandowsky et al., 2017; Stanovich, 2011). At the same time, scepticism is cognitively demanding and effortful (Evans et al., 2013; Kahneman et al., 2021; Kurzban et al., 2013; Normand, 2008; Westbrook et al., 2015), and when cognitive resources are taxed, attentional control decline, promoting reliance on intuitive responses (Hagger et al., 2010).

In high-stakes domains like medicine, law, and finance, professionals must make decision under cognitive load and uncertainty (Kahneman et al., 2009; Shanteau, 1992). Such conditions require epistemic vigilance to assess credibility, detect deception, and resist flawed intuitions (Mercier et al., 2011; Pennycook et al., 2019). This vigilance takes domain-specific forms: clinical scepticism in healthcare (Croskerry, 2003), evidentiary reasoning in law (Vrij et al., 2017), or professional scepticism in auditing (Hurtt, 2010). In auditing, scepticism, also known as professional scepticism, underpins fraud detection, risk assessment, and overall audit quality (Gajewski et al., 2024a; PCAOB, 2022, 2023). It is formally defined as *‘a questioning mind, alertness to misstatement risk, and critical assessment of evidence’* (Gajewski et al., 2024a). Auditors must detect subtle anomalies under tight deadlines and complex information (Asare et al., 2000; Board, 2012; Johari et al., 2019; Lambert et al., 2017; McDaniel, 1990; Otley et al., 1996; Svanström, 2016). Effective professional scepticism requires diagnostic discernment, distinguishing benign from aggressive reporting cues rather than simply rejecting information. Low professional scepticism leads to financial misreporting, undetected fraud, and systemic risk (Hurtt, 2010; Nelson, 2009).

To capture this construct, in this study we used the Phillips (1999) audit task. This task presents realistic evidence items that participants classify as aggressive or non-aggressive, providing a validated behavioral index of skeptical decision-making and judgment. The items in this task are drawn from real-world contexts and expert-validated, capturing the anomaly detection central to fraud identification while maintaining experimental control. Aggressive financial reporting refers to accounting practices that present a company financial position more favourably than justified by the underlying facts or authoritative guidance (Phillips, 1999).

The digitized pace of life has sharply increased information demands, with individuals processing up to 74 GB daily (Bohn et al., 2012). Such volumes exceed cognitive capacity, creating overload (Sweller, 1988), and straining executive functions and attention. The Time-Based Resource-Sharing (TBRS) model holds that executive functions and memory compete for limited resources, making performance vulnerable under load (Barrouillet et al., 2004; Barrouillet et al., 2007; Barrouillet et al., 2012). Overload impairs inhibitory control (Barzykowski et al., 2018; Mazzoni, 2019; Vannucci et al., 2015), problem-solving (Ninomiya et al., 2024; Sweller, 1988), and decision-making (Ordali et al., 2024), while elevating fatigue (Kahneman et al., 2021; Sweller, 1988) and metabolic strain (Backs et al., 1994; Csipo et al., 2021). These effects are acute in tasks requiring vigilance and critical assessment, such as professional scepticism, required for judgement and decision-making. As prefrontal resources are taxed, individuals shift toward heuristic processing that undermines skeptical inquiry (Evans et al., 2005; Gigerenzer, 2008; Van Gestel et al., 2021). In auditing, this shift reduces risk detection, fraud identification, and audit quality (Austin, 2023; Bedeir, 2024; Nelson, 2009; Rose, 2007). Yet, few studies have experimentally examined how cognitive load undermines professional scepticism in judgment and decision-making context, or what mechanisms might help preserve it under constraint.

Given these challenges, default options offer a promising way to reduce the cognitive strain of professional scepticism. Default nudges are pre-set actions that apply unless individuals actively choose otherwise (Thaler & Sunstein, 2008). Though opt-out remains possible, default nudges shape behavior via status quo bias (Gajewski et al., 2022; Meunier et al., 2024b) and have been shown to influence risk-taking, financial choices, and socially responsible investment (Gajewski et al., 2022; Gajewski et al., 2024b; Mertens et al., 2022; Meunier et al., 2024a; Meunier et al., 2024b; Münscher et al., 2016). Their strength lies in reducing cognitive demands: simplifying choices, conserving resources, and sustaining performance under overload (Kool et al., 2018; Marchiori et al., 2017; Meunier et al., 2024a; Sunstein, 2023; Thaler et al., 2008). Neurocognitively, default nudges align with energy-efficient heuristics that balance effort and accuracy (Gigerenzer et al., 2011). Thus, they can function not only as behavioral nudges but as cognitive scaffolds that support effortful processes like professional scepticism when these are most vulnerable (Van Gestel et al., 2021).

### 1.1. Current Study

Although prior work has shown that default nudges can shape behavior under cognitive load, the underlying mechanism remains debated. In the context of dual-processing theory, some accounts suggest that nudges operate primarily by exploiting Type 1 processes, steering intuitive responding when analytic resources are limited (Evans et al., 2013). Others argue that default nudges reduce effort costs, freeing up resources for more deliberate Type 2 reasoning (Kool et al., 2018; Sunstein, 2023). Similarly, models such as TBRS highlight competition between memory and executive functions such as decision-making, but it is unclear whether default nudges compensate for this resource conflict or simply redirect responses heuristically. By testing defaults in a task requiring professional scepticism, a cognitively demanding, deliberative process, we aim to inform this broader debate.

Building on this framework, we ran two online experiments using Phillips’ audit task to test how cognitive load and default nudges interact in shaping professional scepticism. Our contribution is threefold. First, we extend prior work on load and nudges (i.e., (Van Gestel et al., 2021)) by applying it to *professional scepticism in auditing*, a domain where judgment quality has direct societal and financial consequences. Second, we move beyond the general question of whether nudges help under load to examine the *boundary conditions of their effectiveness*, showing that their benefits depend on whether the default aligns with the most probable correct response. Third, we highlight the practical trade-offs of nudge placement, where alignment with majority responses increases efficiency but risks missed fraud cues, while misalignment boosts vigilance but at the cost of false alarms.

Together, these studies contribute to dual-process and decision architecture models by suggesting that default nudges may function as cognitive scaffolds that support judgment under load, and they offer practical strategies for sustaining critical thinking in real-world, high-stakes settings.

## 2. General Materials and Methods

Participants were recruited from Prolific (www.prolific.com), an online study pre-registered subject pool. This platform guarantees the recruitment of a homogenous sample that fits well with the specified inclusion criteria (see Inclusion Criteria for a review). Once the inclusion criteria were established, the study advertisements were exclusively visible to participants meeting these eligibility requirements. Within the advertisement on Prolific, participants were provided with a brief overview of the study and the link to the online experiment. The Gorilla Experiment Builder platform (www.gorilla.sc) (Anwyl-Irvine et al., 2020) was used to build and host the online experiments. All procedures were approved by the local ethics committee of the School of Management ESSCA (Reference No. Avis 7/2024, dated 19 August 2024). All participants provided electronic informed written consent prior to the experiment.

Prior to the experiments hosted on Gorilla, registered participants were informed about the study requirements, including being in a quiet and distraction-free environment. A brief description of the study objectives was provided, along with details on participants’ roles during the experiment, the estimated time required to complete the tasks, the compensation they would receive upon successful completion, and the system requirements (i.e., using Google Chrome on a desktop or laptop computer). It was explicitly stated that participants would not be compensated if they failed to meet the system requirements, did not complete the experiment successfully, or achieved a low-level performance (see Stimuli and Procedure for a review). Participants then were directed to the consent form before reporting their audit experience, which ranged from ‘less than 1 year’ to ‘more than 10 years’ (only presented to participants of experiment 2) (Table 1). Following this, they advanced to the main tasks.

**Table 1.** The table below presents the number (N) and percentage (%) of professional auditors based on their years of experience in auditing, as well as the extent to which the pre-selected option influenced their decision-making-based professional scepticism.

| Category | N | Percentage (%) |
| --- | --- | --- |
| Years of Experience |  |  |
| Less than 1 year | 83 | 31.3 |
| 1–5 years | 136 | 51.3 |
| 5–10 years | 28 | 10.6 |
| More than 10 years | 8 | 3.0 |
| Other | 9 | 3.4 |
| Influence of Pre-Selected Options |  |  |
| Not at all | 17 | 14.9 |
| A little | 22 | 19.3 |
| Somewhat | 29 | 25.4 |
| Quite a bit | 22 | 19.3 |
| A lot | 14 | 12.3 |
| Extremely | 10 | 8.8 |

After completing the main experiment, to check the effectiveness of manipulation, participants who took part in experiment 2 were asked to rate the extent to which they perceived the “pre-selected” options or default nudges in the test as influencing their decisions regarding each piece of evidence. Responses were recorded on a 6-point Likert scale ranging from “not at all (1)” to “extremely (6)” (Table 1). Both experiments ended with a debriefing.

### 2.1. Inclusion Criteria

Inclusion criteria for both experiments were a) being 18 years old or more and under 60 years old, b) having no dyslexia or dyscalculia, c) having normal vision with no color-blindedness, and for experimen 2 d) being either a professional auditor or student of finance or business with knowledge of auditing. Since the primary objective of experiment 1 was to only validate the cognitive load manipulation and domain-specific expertise was not expected to affect performance, non-auditor participants were recruited. The age range was selected to preserve consistency in cognitive functioning (Salthouse, 2009; Salthouse et al., 2008).

### 2.2. Experimental Measures

#### 2.2.1. Dot Memory Task (DMT)

The Dot Memory Task (DMT; (Bethell-Fox et al., 1988)) is a validated (Sliwinski et al., 2018) visuospatial WM paradigm widely used to manipulate and index cognitive load. Participants briefly encode a grid containing a dot pattern, following a distractor task (also known as retention phase), and recall positions of dots in a matrix. Load is varied by matrix/grid size and dot number, which systematically modulates recall difficulty. The task reliably taxes WM resources and has been used in similar prior cognitive load and default nudge research, making it a suitable tool for experimentally inducing and measuring load in decision-making contexts (Van Gestel et al., 2021).

#### 2.2.2. Phillips’ Audit Task

The Phillips’ audit task is a widely used tool in the field of auditing (Phillips, 1999). In this task, participants are presented with background information describing audit scenarios involving clients who manufacture and market water, electricity, and natural gas meters. The information notes that the metering industry has experienced recent growth due to technological innovations and that the client company is a growing entity with increasing revenue, income, and assets. After reading this background information, participants proceed to a review of audit evidence consisting of 45 statements, each related to one of 14 financial statement accounts. We selected a subset of 30 statements from the original 45-item task to reduce task length while maintaining conceptual coverage. Of these, 21 were classified as non-aggressive and 9 as aggressive, based on prior coding in Phillips (1999).

These were distributed equally across three blocks, with 7 non-aggressive and 3 aggressive items per block. For example, an aggressive item might be: *“The principal on the $5,000,000 note receivable will be repaid by Bronzco in increasing annual instalments that do not begin until 2026.”* In contrast, a non-aggressive item could be: *“Depreciation on the main categories of fixed assets was recalculated and appears appropriate.”*

#### 2.2.3. Attention Check

To confirm that participants were attentive throughout the experiment, we incorporated attention checks into the task. In experiment 1, the attention check was operationalized as the proportion of correct responses on both DMT and arithmetic tasks (adjusted accuracy). In Experiment 2, attention checks were embedded within audit-related scenarios which required participants to perform simple arithmetic calculations. For example, one attention check presented the following statement: “*Meter-Tek purchased 300 units in January and sold 150 units in February, leaving a remaining inventory of 200 units*.” Participants were then asked, “*Is this correct?*”. These items served as embedded comprehension probes and were pseudo-randomized across blocks (two per block; six in total), balancing their positions across participants.

Our rationale for using different attention checks across the two experiments was tied to their different objectives. Experiment 1 was designed only to validate the cognitive load manipulation, so attention was appropriately indexed on the DMT + arithmetic task. In experiment 2, by contrast, we did not require trials to be simultaneously correct on both the DMT and the audit attention check questions (ACQs) for four reasons: (1) filtering on DMT correctness would bias the analysis by removing the very trials where load exerted its influence, thereby blinding the effect of load and nudge on decision-making; (2) higher load naturally produces more DMT errors, leading to disproportionate data loss and non-comparable cells; and (3) the construct of interest was sceptical decision-making (under load), not decision-making only when memory recall was perfect.

In line with Prolific guidelines and similar studies, participants who failed to meet the attention check criteria threshold (at least ≥2) were excluded from the analysis (see Participants for a review) (Black et al., 2025; Hauser et al., 2016).

#### 2.2.4. Objective Scepticism

Consistent with (Gajewski et al., 2024a), we tallied objective scepticism as the percentage of aggressive items correctly identified (coded as “1”) within each condition (i.e., low load), based on the three aggressive items presented. This index captures the degree of analytic, deliberative reasoning applied to complex financial evidence, and is conceptually consistent with established models of scepticism in auditing (Hurtt et al., 2013).

### 2.3. Data Analysis

In both experiments, reaction time (RT) (in seconds (s)) and accuracy (in percentage (%)) were computed as dependent measures indexing cognitive load in the DMT and decision-making-based professional scepticism in the Phillips’ audit task.

RTs of DMT during the recall phase were operationalized as the total time elapsed from the presentation of the empty recall matrix to the participants final dot placement. This measure reflects the full temporal span of the recall process, including memory retrieval, spatial decision-making, and motor execution (Sliwinski et al., 2018). Participants were not permitted to modify their responses once the final dot was placed. In the Phillips’ audit task, it was measured from the moment the audit piece appeared on the screen until participants selected their response. RTs were averaged within participants across all trials in each condition (i.e., LL), yielding one RT score per condition per participant.

Accuracy of DMT was scored dichotomously: a score of 1 was assigned when all target dots were placed in their correct grid locations, and 0 otherwise (i.e., partial placement, wrong placement or no placement at all). This stringent scoring approach prioritized complete recall fidelity, which was essential for assessing the effectiveness of the cognitive load manipulation. Similarly, accuracy of Phillips’ audit task was given 1 for correct and 0 for incorrect response. For both DMT and audit, accuracy was calculated for each participant as the proportion of correct responses within each block of 12 trials for DMT and 10 for Phillips’ audit task (excluding 2 attention trials). This yielded three accuracy values per participant (one per block), which were then averaged across participants and converted to percentage accuracy.

We adopted a lenient approach to outliers. Since the goal was to examine the effect of cognitive load, as a function of processing, on pattern retention in WM without introducing time constraints (Hu et al., 2024), timing constraints were not imposed in the task. Thus, participants reconstructed the pattern *at their own pace*. Given that WM was already taxed by the concurrent Phillips’ audit task, they were expected (and instructed) to reconstruct the pattern as quickly as possible to minimize memory lapse. This approach aimed to isolate cognitive load effects as a function of the complexity of the dot pattern while reducing time pressure. Thus, both RTs and accuracy outliers were retained.

RTs naturally vary due to individual differences in cognitive strategies, attention, and prior experience, all of which are integral to learning and memory. Removing long RTs could eliminate meaningful strategic differences in recall, as some participants may take longer to process and reconstruct patterns in externally valid ways (Miller, 1991). Similarly, accuracy variations arise from cognitive factors like momentary lapses in attention rather than random noise. Removing accuracy outliers could exclude participants whose performance, while seemingly suboptimal, remains informative for understanding cognitive processes (Miller, 2023). Extreme RTs and accuracies provide information about individual differences, making their retention essential for a comprehensive analysis of memory recall in an unconstrained temporal context.

Nonetheless, we also report all analyses of experiments 1, 2 with outliers (± 3 SD) in RTs and accuracy removed (see Supplementary Material for a review). RT outliers were identified and removed per participant, while outlier accuracy rates were detected and excluded at the group level across participants, prior to statistical analyses. Subsequently, participants with over 50% missing responses across DMT or audit trials were excluded from analyses. However, no participants met this criterion.

We adopted a dual approach to inference. ANOVAs and non-parametric tests, where assumptions were violated, were conducted as our primary method of hypothesis testing, consistent with preregistration. In parallel as complementary analysis, null hypothesis significance testing was conducted using an alpha level of *p* < 0.05. Bayes Factors (BF₁₀) were computed using noninformative Jeffreys priors on the variance and a Cauchy prior on the standardized effect size (*r* = 0.707), using the *`‘`*BayesFactor*’* package, version 0.9.2+ (Morey et al., 2015) in R version 4.3.1 (Team, 2023). BF₁₀ < 0.3 provide strong evidence in favour of the null hypothesis, whereas BF₁₀ > 3 indicate strong evidence in favour of the alternative hypothesis (Jeffreys, 1961; Rouder et al., 2012; Rouder et al., 2009).

All *p*-values are reported exactly (unless < 0.001). All factorial analyses included interaction terms (i.e., Load × Nudge), tested directly rather than inferred from separate significance tests. RTs were positively skewed; analyses were repeated with log-transformed values and yielded the same conclusions, so we report untransformed RTs for ease of interpretation. Results are presented with means, standard errors, and standardized effect sizes to aid interpretation.

### 2.4. Data availability statements

Data curation, processing and statistical analysis were conducted in MATLAB, unless otherwise specified (MATLAB, 2019). All data and MATLAB analysis code have been made publicly available at the Open Science Framework (OSF). All raw, curated data, analysis code, and research materials are available at [https://osf.io/s4tc8/]. In addition, we preregistered (before data collection) our general research questions, our analysis strategy, our inclusion criteria, and our projected sample size and all are available at [https://osf.io/agmwk]. A video is provided as an example at [www.vimeo.com].

### 2.5. Participants

A total of 299 participants initially took part in the study (38 in experiment 1, 261 in experiment 2). Across both experiments, data from 240 out of 299 participants were included in the analyses, including only those who passed attention checks (see Experimental Measures for a review). The attention check pass rates were 94.7%, and 78.1% for experiments 1 and 2, respectively. These rates are well above the Prolific benchmark for standard ACQs, which is approximately 69% (Peer et al., 2017). This suggests that participant engagement and data quality were high across experiments, particularly given that ACQs are known to be a valid and conservative tool for screening inattentive respondents in online studies. We attribute the relatively large difference in pass rates between the two experiments to the greater complexity of the second experiment.

In accordance with recommendations for transparent reporting of sample size justification (Lakens, 2022), our recruitment strategy combined practical and statistical considerations. First, given the specificity of our target population (i.e., professional auditors), we aimed to recruit the largest feasible sample. Second, we conducted a-*priori* power analyses separately for each experiment. These analyses were based on the smallest effect of theoretical interest in this research context, rather than conventions alone. The selected effect size was informed by prior studies reporting similar magnitudes under comparable conditions (Steffel et al., 2016; Van Gestel et al., 2021). While Cohen (1988) classifies *d* = 0.5 as a “medium” effect, we follow Correll et al. (2020) in grounding our choice in theoretical rationale and empirical precedent. To guard against overestimation, we also considered cautions regarding inflated published effects (Anderson et al., 2017), making our target conservative yet meaningful for professional scepticism in judgement and decision-making. This justification applies across both experiments. Power analyses were conducted using the `‘pwrss`’ package (Bulus, 2023) in R version 4.3.1 (Team, 2023) and effect sizes are reported using eta squared (η²), consistent with the values used in planning.

## 3. Experiment 1: Validation of Cognitive Load Manipulation

In Experiment 1, the objective was to validate the cognitive load manipulation using the DMT, with performance indexed by RT and accuracy. We predicted that increasing grid size and the number of dots would increase memory load, resulting in slower RTs and reduced accuracy. As expected, RTs increased and accuracy declined as load intensified, confirming the effectiveness of the manipulation.

### 3.1. Methods

#### 3.1.1. Participants

Thirty-six participants (out of 38) completed the online experiment (age range: 18 - 60 years, M = 38.4 ± 11.87; 50% female (*N* = 18), 41.66% male (*N* = 15), and 8.33% rather not to say (*N* = 3)). Participants received fair compensation based on Prolific recommended hourly rate (£9 or €10.8 per hour), amounting to €0.75 for approximately 5 mins of participation. An a-priori power analysis indicated that a total sample size of 24 participants would provide 80% power to detect a moderate effect size (*d* = 0.5 or η² = 0.059), assuming an α = 0.05 (Cohen, 1988), using a one-way repeated-measures ANOVA with four load levels (Minimal, Low, Medium and High). As no data had been collected at the planning stage, power analysis relied on parametric assumptions, despite the final use of the non-parametric Friedman test due to normality violations. A total of 36 participants completed the study, exceeding the preregistered minimum and ensuring adequate statistical power.

#### 3.1.2. Tasks and procedure

Figure 1 shows an overview of the experimental design. The DMT (Bethell-Fox et al., 1988) was used to modulate cognitive load by manipulating the complexity of visual WM retention across four conditions (minimal, low, medium, and high loads) (Miyake et al., 2001). The task began with the presentation of a fixation cross for 1s, followed by an encoding period; participants were presented with a cue trial in which the dot pattern appeared in the grid for a pre-determined duration (1s). Following the presentation of the grid, it disappeared, marking the onset of a brief retention interval intended to minimize subvocal rehearsal of the dot positions and encourage reliance on WM. During this interval, participants engaged in a secondary task involving arithmetic (i.e., subtraction), where responses were constructed such that, the correct answer was always ≤ 10. They were required to complete this task within a 5s time window, indicated by a countdown timer displayed in the top-right corner of the screen. Immediate visual feedback was provided after each trial: a red cross for incorrect and a green checkmark for correct responses. In the recall phase, participants reconstructed the dot pattern within a self-paced response window. This task was intentionally designed to be readily manageable for participants while simultaneously imposing cognitive demands to tax WM resources. Participants started with a 3-trial practice block. Participants were required to achieve an accuracy of 100% (3 out of 3 correct responses) before proceeding to the main task; otherwise, the task was repeated until they met this criterion. All participants passed on their first or second attempt. They then completed four experimental blocks, each consisting of 12 randomly presented trials, resulting in a total of 48 trials across the experiment. The four blocks corresponded to varying levels of cognitive load that were defined the number of dots and size of the grid: minimal (2×2 grid with 2 dots), low (3×3 grid with 2 dots), medium (3×3 grid with 3 dots), and high (4×4 grid with 4 dots) loads. The order of blocks (i.e., load conditions) was counterbalanced across participants, and the trials within each block, including DMT and arithmetic task, were presented in randomized order (Figure 1).

**Figure 1.**
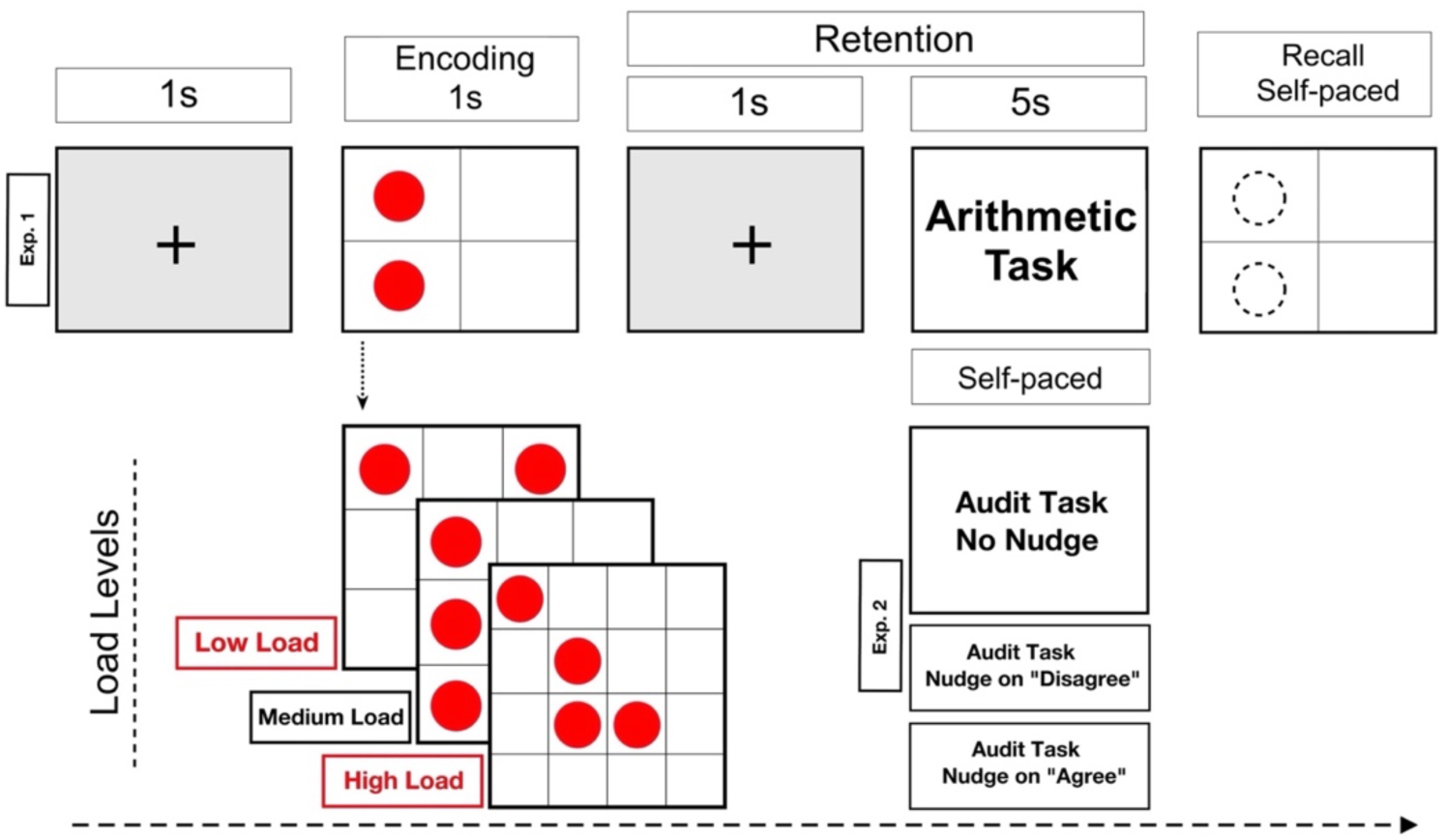
The sequence and timing of the Dot Memory Task (DMT). Each trial began with a fixation cross, followed by an encoding phase (1 s), a retention interval (5 s in experiment 1), and a self-paced recall phase. Experiment 1 included four levels of working memory (WM) load, whereas the main experiment (Experiment 2) adopted the load levels of 2 and 4. The overall structure in experiment 2 was identical to experiment 1, except that the Phillips’ audit task replaced the arithmetic task during retention with no time constraint.

#### 3.1.3. Results: An Increase in Cognitive Load Is Associated with Longer RTs And Reduced Accuracy

Figure 2 shows the behavioural performance of DMT metrics including (i) RTs (Panel 1A) and (ii) adjusted accuracy (Panel 1B) of DMT in experiment 1. First, we calculated ‘adjusted accuracy’, defined as the proportion of trials where participants provided correct responses on both the DMT and the concurrent arithmetic task. Next, we computed a broader (iii) accuracy metric (‘accuracy + math’), which included all correct DMT responses regardless of correctness on the arithmetic task (incl. correct and incorrect responses) (Panel 1C). Finally, we analysed (D) overall DMT accuracy without consideration of performance on the arithmetic task (‘overall accuracy’) (incl. miss, correct and incorrect responses) (Panel 1C). Patterns of results were consistent across all three metrics: performance declined with increasing cognitive load, supporting the effectiveness of the load manipulation.

**Figure 2.**
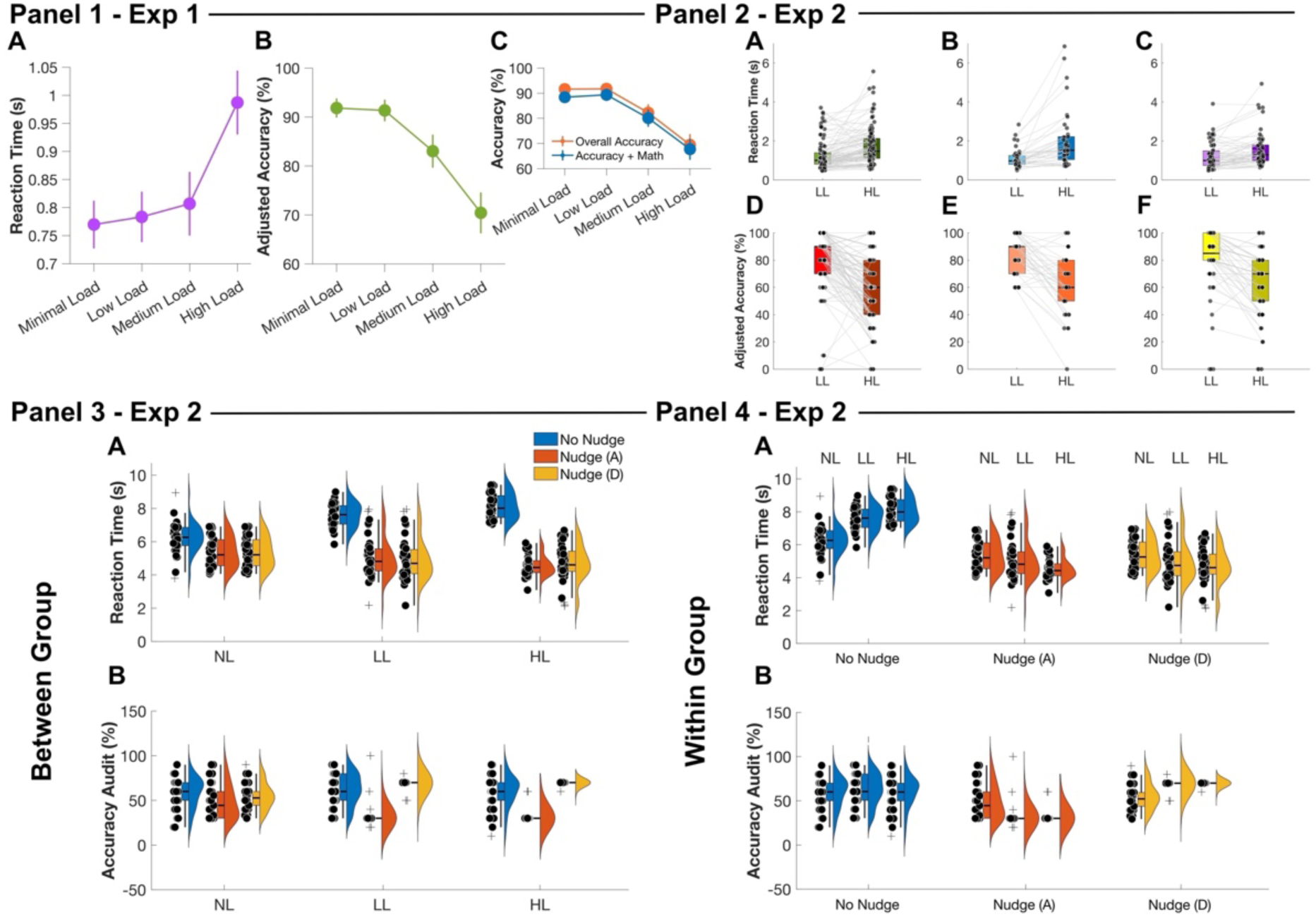
Panel 1A illustrates RTs of DMT, panel 1B depicts adjusted accuracy. Panel 1C shows overall accuracy and accuracy + math. The error bars represent ±1 SEM. Panels 2A, B, and C illustrate the RTs in the DMT per experimental group including no nudge, nudge (A) and nudge (D), and panels 2D, E, and F show the accuracy of the DMT in experimental groups. Panel 3A depicts the behavioural performance on the Phillips’ audit task, showing RTs (s), and 3B show accuracy rate (%) between the experimental groups. Panel 4A displays within-group differences of RTs and accuracy on the Phillips’ audit task. Each whisker represents the interquartile range (IQR), the horizontal black line denotes the mean. The kernel density plots visualize the data distribution. All circles represent individual data points. Abbreviations: NL (No Load), LL (Low Load), and HL (High Load) on x-axes.

A Friedman test revealed significant differences in RT across load conditions (χ^2^(3) = 33, *p* < 0.001, BF_10_ = 7.45) (Panel 1A). Post-hoc Wilcoxon signed-rank tests with Bonferroni correction indicated that RTs under high load were significantly longer than under minimal (Z = −4.59, *p* < 0.001, BF_10_ = 5.27), low (Z = −4.21, *p* < 0.001, BF_10_ = 9.45), and medium (Z = −4.14, *p* < 0.001, BF_10_ = 4.48) loads. However, the minimal and low loads (Z = −0.92, *p* = 0.36, BF_10_ = 0.18), minimal and medium loads (Z = −0.97, *p* = 0.33, BF_10_ = 0.27), and low and medium loads (Z = −1.05, *p* = 0.29, BF_10_ = 0.23) did not show any difference (6 pairwise comparisons).

For adjusted accuracy (Panel 1B), a similar trend was observed. A Friedman test showed significant differences across conditions, (χ²(3) = 25.58, *p* < 0.001, BF_10_ = 15.63). Wilcoxon post-hoc indicated that accuracy under high load was significantly lower than minimal (Z = 4.00, *p* < 0.001, BF_10_ = 15.27), low (Z = 4.47, *p* < 0.001, BF_10_ = 19.45), and medium (Z = 2.71, *p* < 0.001, BF_10_ = 14.48) loads. However, the differences between minimal and low loads (Z = 0.21, *p* = 0.99, BF_10_= 0.18), minimal and medium loads (Z = 2.31, *p* = 0.12, BF_10_= 0.2), and low and medium loads (Z = 2.23, *p* = 0.15, BF_10_= 0.23) were not significant (6 pairwise comparisons). Analysis of accuracy + math (Panel 1C) also showed significant group differences, (χ²(3) = 27.4, *p* < 0.001, BF_10_ = 28.52), with lower performance under high load compared to minimal (Z = 3.88, *p* < 0.001, BF_10_ = 15.47), low (Z = 4.54, *p* < 0.001, BF_10_ = 18.48) loads. However, no significant differences were observed between high and medium loads (Z = 2.61, *p* = 0.053, BF_10_ = 0.28), medium and minimal loads (Z = 2.35, *p* = 0.11, BF_10_ = 0.22), medium and low loads (Z = 2.39, *p* = 0.10, BF_10_ = 0.18), or minimal and low loads (Z = −0.22, *p* = 0.99, BF_10_ = 0.19) (6 pairwise comparisons). Finally, for overall accuracy (Panel 1C), a significant effect of load emerged, (χ²(3) = 28.00, *p* < 0.001, BF_10_ = 28.1). Accuracy under high load was again significantly lower than minimal (Z = 4.03, *p* < 0.001, BF_10_ = 20.51), low (Z = 4.58, *p* < 0.001, BF_10_ = 17.46), and medium loads (Z = 2.72, *p* < 0.001, BF_10_ = 15.1). But there was no significant difference between minimal and low loads (Z = −0.08, *p* = 0.99, BF_10_ = 0.18), minimal and medium loads (Z = 2.53, *p* = 0.07, BF_10_ = 0.12) and low and medium loads (Z = 2.44, *p* = 0.09, BF_10_ = 0.66) (6 pairwise comparisons). These results confirm that the load manipulation was effective, yielding clear behavioural modulation across load levels. Based on these findings, the low and high load conditions were selected for the subsequent experiments.

## 4. Experiment 2 (main): Effect of Cognitive Load on Phillips’ Audit Task and its Interaction with Default Nudges

In Experiment 2, the objectives were to examine (A) whether cognitive load influences performance on the Phillips’ audit task, (B) how default nudges interact with load to shape professional scepticism in judgement and decision-making, and (C) if the placement of nudges (agree vs. disagree) affects their effectiveness. Participants were randomly assigned to one of the three groups: no nudge, nudge (agree (A)), and nudge (disagree (D)) and each completed tasks under No Load (NL), Low Load (LL), or High Load (HL) conditions, adapted from the DMT manipulation in Experiment 1. We predicted that higher load would impair performance, reducing accuracy and slowing RTs, and that default nudges would improve efficiency through faster RTs and greater accuracy. Results showed that increasing load reliably slowed RTs but did not significantly reduce accuracy, suggesting depletion primarily affects efficiency rather than accuracy. Default nudges reduced RTs across groups but influenced accuracy selectively: in the nudge (A) group, accuracy decreased, whereas in the nudge (D) group, accuracy improved.

### 4.1. Methods

#### 4.1.1. Participants

All experiment 2 groups were recruited and run during the same data-collection window and analysed as a single mixed design. The final analysis included data from 204 participants out of 261 (age range: 18 - 60 years, M = 30.06 ± 9.2 years; 46.29% female (*N* = 91), 39.81% male (*N* = 129), 13.88% rather not to say (*N* = 1)), 108 out of 137 in no nudge, 38 out of 51 in the nudge (D), and 58 out of 73 in the nudge (A) groups.

A-priori power analysis showed that a minimum sample of 28 participants would provide 80% power to detect a moderate effect size (η² = 0.059) with *α* = 0.05 for the main effect of load (within-subjects). For detecting between-subjects main effects or interaction effects in the three-group mixed factorial design, the required total sample sizes were *N* = 106 and *N* = 35, respectively, assuming *α* = 0.05 and 80% power. The between-subjects factor had three levels (no nudge, nudge (D), and nudge (A)) and the within-subjects factor had three levels of cognitive load (NL, LL, and HL).

Participants received €2.15 for approximately 15 mins of participation. To further promote engagement and task performance, participants were also offered a €0.05 bonus for each correct response on both the DMT and Phillips’ audit task.

#### 4.1.2. Tasks and procedure

The main experiment replicated the structure of the prior experimental design with a key modification to integrate Phillips’ audit task instead of the arithmetic (Figure 1). Additionally, we employed NL condition, alongside the LL and HL conditions from experiment 1. The NL condition excluded the DMT, such that participants completed only the Phillips’ audit task. The reason for including the NL condition was twofold: first, to establish a baseline for evaluating the impact of cognitive load; and second, to simulate real-world decision-making contexts in which individuals operate without concurrent memory demands.

Prior to the task, participants were presented with a detailed audit scenario, which served as the context for evaluating subsequent audit pieces of evidence. The task began with a fixation cross, followed by the encoding of a randomly generated pattern of dots, and was then followed by a retention interval. During designated intervals within the experiment, participants were presented with individual audit pieces of evidence sequentially. For each piece of evidence, they were required to make a judgment based on the initial audit scenario, answering the question: *“Is this piece indicative of aggressive reporting?”* Participants in the no nudge group responded at their own pace by selecting either “Yes” or “No.” In the NL condition, participants viewed the evidence and responded immediately after the first fixation cross, with no DMT included. Importantly, in the no nudge group, participants always made an active choice between “Yes” and “No,” with no default pre-selected. Thus, the no nudge group represents a true forced-choice baseline.

In the nudge (D) or nudge (A) groups, however, a default nudge was incorporated, where instead of actively selecting “Yes” or “No”, a response was pre-selected for the participants. Specifically, two options were provided: “Agree” and “Disagree,” with the pre-selected response set to “Disagree” in nudge (D), reflecting its higher likelihood of being correct (∼70% of audit items were non-aggressive, necessitating “Disagree” as the correct response). Participants were instructed to carefully read the audit pieces: if they disagreed that the piece was indicative of aggressive auditing (i.e., agreed with the pre-selected response), they simply clicked “Continue.” However, if they judged the statement to be aggressive, they were required to drag the checkmark from “Disagree” to “Agree” before proceeding. In the nudge (A) condition, the pre-selected option was “Agree.” The response mapping was intentionally asymmetric: accepting the pre-selected default required a single click, whereas overriding it required two actions (dragging the checkmark and clicking “Continue”). This design increased the effort cost of rejecting the default, thereby making adherence to the pre-selected option less effortful than deviation.

The placement of the default nudge was fixed across all trials of nudge (D) (always on “Disagree”) and nudge (A) (always on “Agree”).

#### 4.1.3. Results: Default Nudges Enhance Response Efficiency by Decreasing RTs and Improving Accuracy Under Cognitive Load

We first tested the effect of cognitive load on WM using the DMT in all three groups. Rank RTs were computed for each participant in the ‘LL’ and ‘HL’ conditions and compared using a Wilcoxon signed-rank test, which revealed significantly slower responses under HL (Z = −4.533, *p* < 0.001, BF_10_ = 25.4; Panel 2A) relative to LL in no nudge, (Z = −4.793, *p* < 0.001, BF_10_ = 19.67; Panel 2B) in nudge (A), and (Z = −4.533, *p* < 0.001, BF_10_ = 24.65; Panel 2C) in nudge (D) groups. Accuracy showed a corresponding drop under HL (Z = 5.089, *p* < 0.001, BF_10_ = 31.22; Panel 2D) in no nudge, (Z = 4.32, *p* < 0.001, BF_10_ = 15.4; Panel 2E) in nudge (A) and (Z = 5.089, *p* < 0.001, BF_10_ = 19.6; Panel 2F) and in nudge (D) groups. These results replicate the cognitive load manipulation from experiment 1 and validate that the DMT successfully imposed varying levels of WM load in all groups.

To test the effect of default nudges and load on Phillips’ audit task performance, a two-way mixed ANOVA with Greenhouse-Geisser correction was conducted. The between-subjects factor was nudge condition (no nudge, nudge (D), nudge (A)), and the within-subjects factor was cognitive load (NL, LL, and HL). For Phillips’ audit task RTs, there was a significant main effect of nudge condition (F (2, 201) = 392.65, *p* < 0.001, η² = 0.77, BF_10_ = 5.45) but no significant main effect of load (F (1.87, 402) = 1.31, *p* = 0.27, η² = 0.006, BF_10_ = 0.26). A significant nudge × load interaction emerged (F (3.74, 402) = 52.36, *p* < 0.001, η² = 0.34, BF_10_ = 6.36) (Panel 3A). For accuracy, the main effect of nudge condition was significant (F (2, 201) = 84.26, *p* < 0.001, η² = 0.46, BF_10_ = 5.63), while the main effect of load was not (F (1.84, 402) = 0.423, *p* = 0.63, η² = 0.002, BF_10_ = 0.29). The nudge × load interaction was also significant (F (3.69, 402) = 22.9, *p* < 0.001, η² = 0.19, BF_10_ = 7.74; Panel 3B), suggesting that the impact of nudges on accuracy was conditional on the level of cognitive load.

For RTs at all load levels, no nudge vs. nudge (A), and no nudge vs. nudge (D) were significant (NL; (MD = 0.068, 95% CI= [0.03, 0.11], *p* < 0.001, BF_10_ = 5.67) and (MD = 0.08, 95% CI= [0.03, 0.12], *p* < 0.001, BF_10_ = 6.02)), (LL; (MD = 0.25, 95% CI= [0.21, 0.29], *p* < 0.001, BF_10_ = 9.71) and (MD = 0.25, 95% CI= [0.21, 0.3], *p* < 0.001, BF_10_ = 7.07)), and (HL; (MD = 0.36, 95% CI= [0.32, 0.39], *p* < 0.001, BF_10_ = 4.9) and (MD = 0.34, 95% CI= [0.3, 0.38], *p* < 0.001, BF_10_ = 4.43)). Whereas nudge (A) vs. nudge (D) were not significant across all load levels, ((NL; (MD = 0.008, 95% CI= [−0.04, 0.061], *p* = 0.99, BF_10_ = 0.03)), (LL; (MD = 0.006, 95% CI= [−0.04, 0.06], *p* = 0.99, BF_10_ = 0.07)), and (HL; (MD = −0.01, 95% CI= [−0.059, 0.032], *p* = 0.99, BF_10_ = 0.09)) (9 pairwise comparisons).

For accuracy, at NL, only no nudge vs. nudge (A) was significant (MD = 8.85, 95% CI= [2.25, 15.44], *p* = 0.004, BF_10_ = 3.76), but no nudge vs. nudge (D) and nudge (A) vs. nudge (D) were not significant (MD = 2.089, 95% CI= [−5.55, 9.73], *p* = 0.99, BF_10_ = 0.06) (MD = −6.76, 95% CI= [−15.22, 1.69], *p* = 0.16, BF_10_ = 0.44). At LL, no nudge vs. nudge (A), no nudge vs. nudge (D) and nudge (A) vs. nudge (D) were all significant (MD = 26.78, 95% CI= [20.51, 33.05], *p* < 0.001, BF_10_ = 6.72), (MD = −10.88, 95% CI= [−18.14, −3.61], *p* = 0.001, BF_10_ = 6.09), and (MD = −37.66, 95% CI= [−45.69, −29.62], *p* < 0.001, BF_10_ = 6.67). Similarly, at HL, all comparisons were significant including no nudge vs. nudge (A) (MD = 25.54, 95% CI= [19.88, 31.2], *p* < 0.001, BF_10_ = 6.74), no nudge vs. nudge (D) (MD = −13.16, 95% CI= [−19.72, −6.61], *p* < 0.001, BF_10_ = 7.04), nudge (A) vs. nudge (D) (MD = −38.7, 95% CI= [−45.96, −31.45], *p* < 0.001, BF_10_ = 6) (9 pairwise comparisons).

These findings demonstrate that default nudges consistently enhance response speed, regardless of their placement. However, their impact on accuracy is load-dependent and context-sensitive: nudges aligned with the most likely correct response (nudge (D)) bolster performance under load, whereas misaligned nudges (nudge (A)) can impair accuracy.

#### 4.1.4. Results: Default Nudges Placement Shifts the Balance Between Detecting Aggressive Items and Avoiding False Alarm

As a complementary analysis, we examined how default nudge placement influences objective scepticism across cognitive load conditions, Kruskal-Wallis tests revealed significant group differences at all load levels: (NL; χ² (2) = 40.41, *p* < 0.001, BF_10_ = 25.11, LL; χ² (2) = 52.67, *p* < 0.001, BF_10_ = 36.08, and HL; χ² (2) = 44.49, *p* < 0.001, BF_10_ = 30.12). To further investigate these effects, Mann-Whitney U tests were performed for pairwise comparisons with adjusted *p-*values. In the NL condition, significant differences were observed between no nudge and nudge (D) (*p* = 0.01, BF_10_ = 7), no nudge and nudge (A) (*p* < 0.001, BF_10_ = 8.44), and nudge (D) and nudge (A) (*p* < 0.001, BF_10_ = 7.41). In the LL condition, significant differences emerged between no nudge and nudge (D) (*p* < 0.001, BF_10_ = 5.18), no nudge and nudge (A) (*p* < 0.001, BF_10_ = 5.72), and nudge (D) and nudge (A) (*p* < 0.001, BF_10_ = 6.2). In the HL condition, performance accuracy also significantly differed between no nudge and nudge (D) (*p* < 0.001, BF_10_ = 5.24), no nudge and nudge (A) (*p* < 0.001, BF_10_ = 7.09), and nudge (D) and nudge (A) (*p* < 0.001, BF_10_ = 6.13). Taken together, these results indicate that when default nudges align with the most probable response, overall, Phillips’ audit task accuracy improves. On the other hand, nudging toward less probable response appears to increase sensitivity to aggressive reporting, enhancing scepticism at the cost of more classification errors (Figure 3, Panel A).

**Figure 3.**
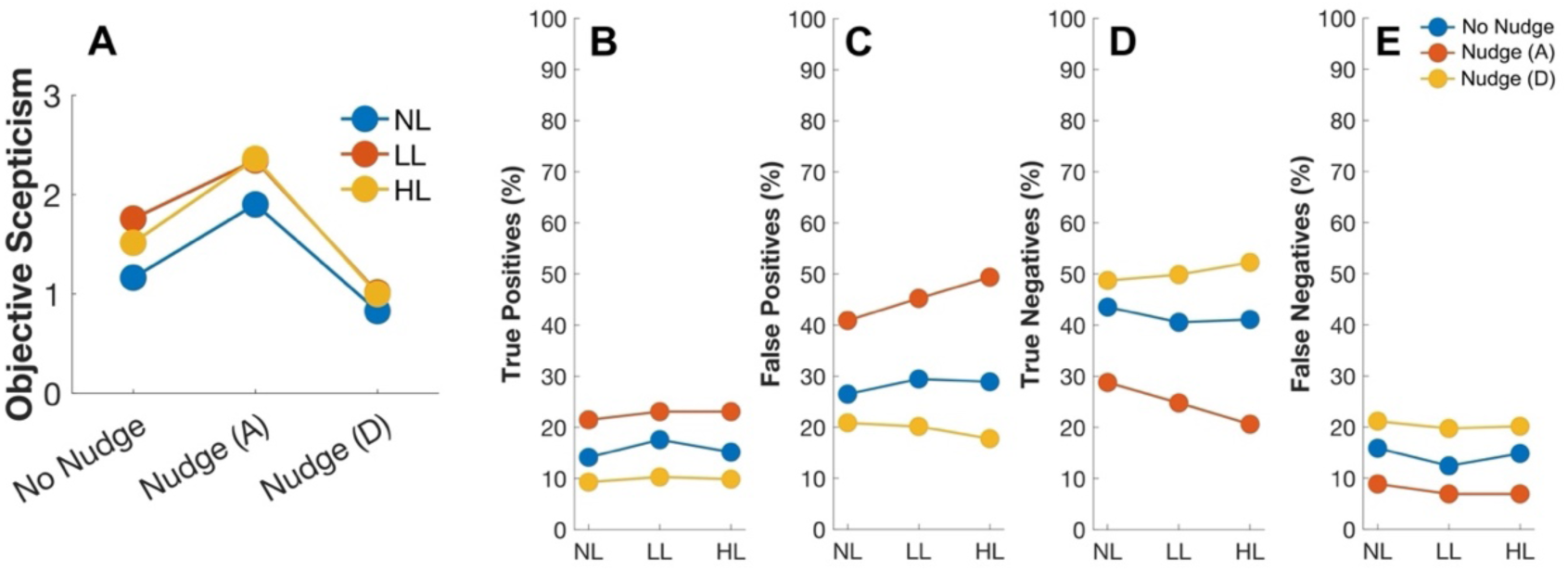
Panel A shows objective scepticism across the no nudge, nudge (A), and nudge (D) groups under NL, LL, and HL conditions. Panels B, C, D, and E display the objective scepticism across nudges conditions for each outcome type: true positive (Panel B), false positive (Panel C), true negative (Panel D), and false negative (Panel E).

To further assess the effect of default nudges placement on the correct detection of aggressive and non-aggressive items, we analysed the general response trends across all three experimental groups (no nudge, nudge (A), and nudge (D)) (Figure 3). The Kruskal-Wallis test revealed significant differences across cognitive load conditions for all response types: true positives (TP; χ²(2) = 138.262, *p* < 0.001, BF_10_ = 18.5), false positives (FP; χ²(2) = 113.559, *p* < 0.001, BF_10_ = 16.83), false negatives (FN; χ²(2) = 138.1, *p* < 0.001, BF_10_ = 17.19), and true negatives (TN; χ²(2) = 112.771, *p* = 0.001, BF_10_ = 16.7). Pairwise comparisons using the Mann-Whitney U test, with Bonferroni-adjusted p-values, indicated that TP rates differed significantly between no nudge and nudge (D) (*p* < 0.001, BF_10_ = 7.13), no nudge and nudge (A) (*p* < 0.001, BF_10_ = 5.67), and nudge (D) and nudge (A) (*p* < 0.001, BF_10_ = 5.04). A similar pattern emerged with the same order (i.e., no nudge and nudge (D) and so on) for FP (BF_10_ = 6.27, BF_10_ = 9.13, BF_10_ = 4.89), FN (BF_10_ = 7.54, BF_10_ = 5.32, BF_10_ = 10), and TN (BF_10_ = 8.41, BF_10_ = 4.12, BF_10_ = 6.78) rates, with all pairwise comparisons yielding significant adjusted *p*-values (*p* < 0.001). These results suggest that nudging participants toward a response with lower probability of correctness (i.e., “Agree”) increases their ability to detect aggressive items (higher hit rate; Panel B). However, this placement also leads to more false alarms, as participants are more likely to misclassify non-aggressive items as aggressive (Panel C). In contrast, nudges placed on the response with higher prior probability of correctness (i.e., “Disagree”) improve the accurate identification of non-aggressive items (higher correct rejection rate; Panel D), but reduce the detection of aggressive items, resulting in more misses (i.e., aggressive items judged as non-aggressive; Panel D).

## 5. General Discussion

In this online study, we investigated whether default nudges influence professional scepticism under varying levels of cognitive load during audit judgment. Experiment 1 validated the effectiveness of our WM load manipulation, showing that HL impaired accuracy and increased RTs relative to minimal, low, and medium load conditions. Based on this behavioural differentiation, we selected the LL and HL conditions for subsequent experiment.

Experiment 2 demonstrated how default nudges interact with cognitive load to influence audit performance. Default nudges reliably reduced RTs, and their effect on accuracy emerged only under heightened cognitive load, indicating that nudges may act as compensatory tools in demanding contexts. In addition, we tested the contextual sensitivity of default nudges, showing that their effectiveness depends on their alignment with the most probable response. Default nudges facilitated faster and more accurate performance when aligned with high-probability responses (i.e., “Disagree”), but undermined accuracy when misaligned (i.e., “Agree”). This asymmetry is critical to the intervention: default nudges provide a comfort anchor that reduces the need for effortful responding under load.

### 5.1. Cognitive Load Modulates Both the Speed and Accuracy of the WM Task

Our findings show that manipulating cognitive load significantly modulates RTs and impedes accuracy, both of which are key indicators of WM function. Elevated cognitive load resulted in prolonged RTs and reduced accuracy, reflecting load-dependent constraints on WM capacity, consistent with the notion that increased intrinsic load or task complexity depletes available WM resources (Sweller, 1988). These results are in line with previous evidence showing that higher memory load disrupts accuracy and slows responses, highlighting inherent fundamental capacity limitations within cognitive processing (Barrouillet et al., 2004; Barrouillet et al., 2007; Camos et al., 2015; Leung et al., 2004; Simon et al., 1963; Zhang et al., 2011; Zhao et al., 2010).

In spatial WM tasks, excessive load impairs both the maintenance, the ability to hold spatial location information in mind over time, and the manipulation, which involves updating or rearranging these locations to make decisions or solve problems, regarding task-relevant information (Leung et al., 2004). Similarly, in response selection paradigms, such as the Simon effect, increasing verbal WM load abolishes the facilitation typically observed for congruent responses, highlighting WM role in conflict resolution (Zhao et al., 2010). The Simon effect refers to the phenomenon where people respond faster and more accurately when the spatial location of a stimulus matches the location of the required response (congruent), even when the stimulus location is irrelevant to the task (Simon et al., 1963). Under low cognitive load, participants exhibit faster responses when stimulus and response locations align, however, this advantage diminishes with rising load, suggesting that higher cognitive demands impair efficient response selection. Furthermore, cognitive load modulates attentional control, with lower loads permitting more effective attentional allocation, while higher loads disrupt attentional guidance (Zhang et al., 2011). Similarly, studies on delayed recall further support this load-dependent interference, as increased concurrent task demands impair both immediate and delayed recall, highlighting the vulnerability of maintenance and retrieval processes to heightened cognitive demands (Camos et al., 2015).

### 5.2. Default Nudges Promote Decision-Making-Based Professional Scepticism Under Cognitive Load Provided That It Is Positioned Correctly

We observed that the implementation of default nudges did not improve the accuracy of professional scepticism performance, but instead only increased response speed when comparing the no nudge and nudge conditions in the absence of cognitive load. Because individuals were not constrained by cognitive load, the availability of cognitive resources likely enabled them to analyse audit information more deliberately, even when this resulted in conscious errors. This supports the view that default nudges primarily function by reducing the cognitive load associated with complex tasks (Meunier et al., 2024a). This finding holds practical relevance for environments such as financial auditing, emergency triage, or regulatory compliance, domains where cognitive overload is common and default nudges might serve as decision aids. In this regard, Van Gestel et al. (2021) state that when individuals have the opportunity to engage in effortful deliberation, they can override default nudges through extensive reasoning. This implies that default effects are robust, but not necessarily impregnable as deliberation may operate in parallel and impact choice simultaneously.

However, under cognitive load, a strikingly different pattern emerged. When the default nudges aligned with the responses with high probability of selection, not only RTs but decision accuracy also improved relative to the no nudge condition. But when the default nudges were placed on the less probable response, accuracy significantly declined. There are two mutually inclusive possibilities for this. The first possibility suggests that 1) individuals frequently shift processing strategies under cognitive load, often relying on simple heuristics, which can preserve or even improve decision accuracy in certain contexts. This adaptive strategy may help explain the success observed in Phillips’ audit task performance (Brunye et al., 2018; Gigerenzer et al., 2011; Sisakhti et al., 2021). This suggests that under cognitive load, individuals show a strong propensity to adhere to the default option, regardless of its alignment with the optimal choice. Further, this highlights the critical role of default nudges architecture in ensuring its effectiveness, particularly when cognitive resources are taxed. The second possibility is that 2) under Type 1 processing, default nudges become more effective no matter where they lead individuals to or even, guarantee success. That is, as default nudges take advantage of the heuristics and biases that are correlated with Type 1 processing, they are more effective in exactly those circumstances where people are most likely to use Type 1 processing. Therefore, inhibiting Type 2 reasoning, for example, by time pressure (Evans et al., 2005) or concurrent WM load through more information (Bago et al., 2017) could in theory increase the effectiveness of default nudges.

However, the presence of default nudges did not alter the performance accuracy in HL relative to LL. These findings converge with those of Van Gestel et al. (2021), who demonstrated that the efficacy of the default nudges remains invariant across cognitive load conditions (LL vs. HL). In their study, despite participants experiencing greater difficulty in recalling the dot pattern under HL (demonstrated through RTs), the default nudges effect remained statistically equivalent. They proposed that default nudges are most effective when Type 2 processing is inhibited. This conforms with the default-interventionist model of dual processing (Evans et al., 2013), which contends that Type 1 processing constitutes the baseline mode of cognition, operating autonomously unless actively overridden by Type 2 reasoning. Crucially, the mere availability of cognitive resources (i.e., in LL condition) does not necessitate the individuals’ higher engagement in the task at hand (Van Gestel et al., 2021).

Our results also contribute to ongoing debates in judgment and decision-making theory. The finding that aligned default nudges improved performance only under load suggests that default nudges may function less as prompts for deliberate reasoning and more as heuristics that become attractive when Type 2 resources are constrained (Evans et al., 2013; Gigerenzer et al., 2011). Conversely, the decline in accuracy with misaligned defaults indicates that defaults do not simply conserve cognitive resources but can also exploit intuitive responding (Bago et al., 2017; Evans et al., 2005). This pattern converges with Van Gestel et al. (2021), who showed that defaults are robust under load, supporting the view that default nudges primarily interact with Type 1 processing rather than enhancing Type 2 engagement, particularly in the context of scepticism-based audit judgments. This sheds light on the importance of contextual design in default nudge interventions across different domains unless the task structure actively engages analytic processing, even cognitively capable individuals may default to less effortful choices.

### 5.3. Strategic Placement of Default Nudges Can Serve Different Purposes

Our results show that defaults aligned with the higher-probability response enhanced performance across both low and high load conditions, improving accuracy and reducing RTs. Building on the role of Type 1 processing outlined above (Evans et al., 2013), this efficiency gain has important implications for applied judgement and decision-making. It is particularly valuable in complex environments where information overload can undermine deliberative reasoning (Gigerenzer et al., 2011; Payne et al., 1993).

However, the placement of default nudges on such responses is not without trade-offs. While it increased true negative rates and reduced false positives, thereby improving accuracy in non-aggressive classifications, it also increased false negatives, i.e., instances in which aggressive audit items were missed. This pattern reveals a conservative bias induced by the default: auditors become more likely to follow the expected norm, potentially overlooking critical red flags (Jachimowicz et al., 2019). In practice, this could translate into missed detections of material misstatements or fraud, posing legal and reputational risks for firms. Therefore, while such nudges configuration optimizes average performance, it may inadvertently downplay low-probability but high-cost errors.

In contrast, placing the default nudges on the less probable response induced a more sceptical decision pattern. This configuration significantly increased the detection of aggressive items (true positives) while reducing false negatives, suggesting that it promotes auditors’ vigilance in identifying problematic cases. Importantly, this vigilance came at the cost of increased false positives, and non-aggressive items incorrectly flagged as aggressive, which could contribute to unnecessary escalation, audit inefficiencies, or strained client relationships. Yet from a risk management perspective, such over-detection may be desirable in contexts where the cost of a false negative (i.e., missing a fraud indicator) far outweighs the cost of a false positive (i.e., reviewing a benign case). These findings are consistent with research suggesting that when error costs are asymmetrical, nudging individuals toward the less likely, but more cautious, response can serve as a protective mechanism against catastrophic failures (Arkes et al., 1986; Gigerenzer et al., 2003).

Thus, the decision to place default nudges on the responses with high or low probability of selection should be informed by the specific error-cost asymmetries of the task domain. In domains like auditing and medicine, where the consequences of false negatives can be severe, legal liability, patient harm, regulatory sanction, and thus default nudges on less probable, risk-sensitive options may operate as a safeguard. On the other hand, in settings where efficiency and throughput are critical, and the cost of false positives is more salient, default nudges aligned with the more probable response may optimize decision fluency and minimize unnecessary friction. Although the observed effects of default nudges on decision accuracy were small to moderate in magnitude, their practical impact should not be underestimated. Default nudges are inexpensive, easy to implement, and require no changes to user training or task procedures. In high-stakes domains where even marginal improvements can prevent costly errors, and where decisions are made repeatedly or at scale, such subtle interventions can yield meaningful cumulative benefits. Crucially, our findings suggest that default nudges are not neutral tools; their effectiveness and ethical soundness depend on how well their configuration fits with domain-specific risk profiles and human cognitive constraints (Reijula et al., 2022; Thaler et al., 2008).

While our experiments focused on auditing decisions, the implications of nudge placement extend to other high-stakes domains involving professional judgment under cognitive constraints. Future studies could examine how default configuration interacts with task complexity and cognitive load in fields such as healthcare, legal reasoning, or safety-critical operations, where the cost of judgment errors varies asymmetrically.

## 6. Limitations

First, although the Phillips’ audit task provides an adequate proxy for professional scepticism in judgement and decision-making context, and we recruited participants with auditing backgrounds, online experiments cannot fully capture the high-pressure, high-stakes conditions of professional audit practice. Second, our operationalization of professional scepticism relied on classification accuracy from a relatively small set of aggressive items, which constrains measurement reliability. Third, because analyses averaged across items and treated them as fixed effects, our conclusions apply most directly to the specific tasks used here. These factors warrant caution in generalizing the findings to broader real-world contexts. Nevertheless, the dot memory task and Phillips’ audit task are validated and externally valid measures, respectively, making them reasonable proxies for cognitive load and professional scepticism.

## 7. Conclusion

This study highlights the complex relationship between default nudges and decision-making-based professional scepticism under cognitive load. Our findings demonstrate that cognitive load impairs auditors’ judgment and decision-making efficiency; however, default nudges can mitigate these effects, enhancing both response speed and accuracy, but only when cognitive resources are taxed. Crucially, the effectiveness of default nudges depends on their compatibility with pre-existing response tendencies, with scepticism-based accuracy improving when default nudges reinforce responses with higher probabilities of selection that is not without caveats. These findings highlight the potential for cognitively informed choice architectures to support professional decision-making in high-stakes environments. Future work should explore the long-term impact of such interventions and evaluate their effectiveness in applied auditing contexts.

## Biography of authors

Dr. Mercede Erfanian is currently a Research Associate at ESSCA Lyon. She has a background in cognitive neuroscience and behavioural psychology.

Prof. Luc Meunier is a full professor at ESSCA Aix-en-Provence, and he is an expert in behavioural finance and neurofinance.

Prof. Jean-François Gajewski is a full professor of Finance at iaelyon School of Management and Executive President of the French Finance Association (AFFI). His research focuses on information and financial markets, behavioural and neurofinance, exploring how managers and investors make financial decisions and process information.

## Acknowledgements

The [REMOTAUDIT](https://anr.fr/Project-ANR-21-CE26-0012) project has received funding from the French Research Agency (Agence Nationale de la Recherche (ANR)), grant ANR-21-CE26-0012-02. We are thankful to the ANR team members. We also thank Ehsan Eqlimi for helping with analysis code.

## Supplementary Material

The following section presents results addressing the primary research questions. Reported findings are based on data after excluding outliers that exceeded ±3 SD from the mean. Outlier RTs were identified and removed at the individual participant level, while outlier accuracy scores were detected and excluded at the group level across participants, prior to statistical analyses.

### Experiment 1

As shown in Figure 1, Wilcoxon signed-rank tests revealed that increasing cognitive load from LL to HL significantly impaired performance across all no nudge, nudge (A), and nudge (D) groups. In the no nudge group, RTs were slower under HL compared to LL (Z = –6.429, *p* < 0.001), and accuracy decreased (Z = 7.532, *p* < 0.001). For the nudge (A) group, HL also produced slower RTs (Z = –4.533, *p* < 0.001) and lower accuracy (Z = 5.089, *p* < 0.001) relative to LL. Similarly, in the nudge (D) group, HL yielded slower RTs (Z = –4.793, *p* < 0.001) and reduced accuracy (Z = 4.323, *p* < 0.001) compared to LL (Figure, Panel 1A, B).

**Figure 1.**
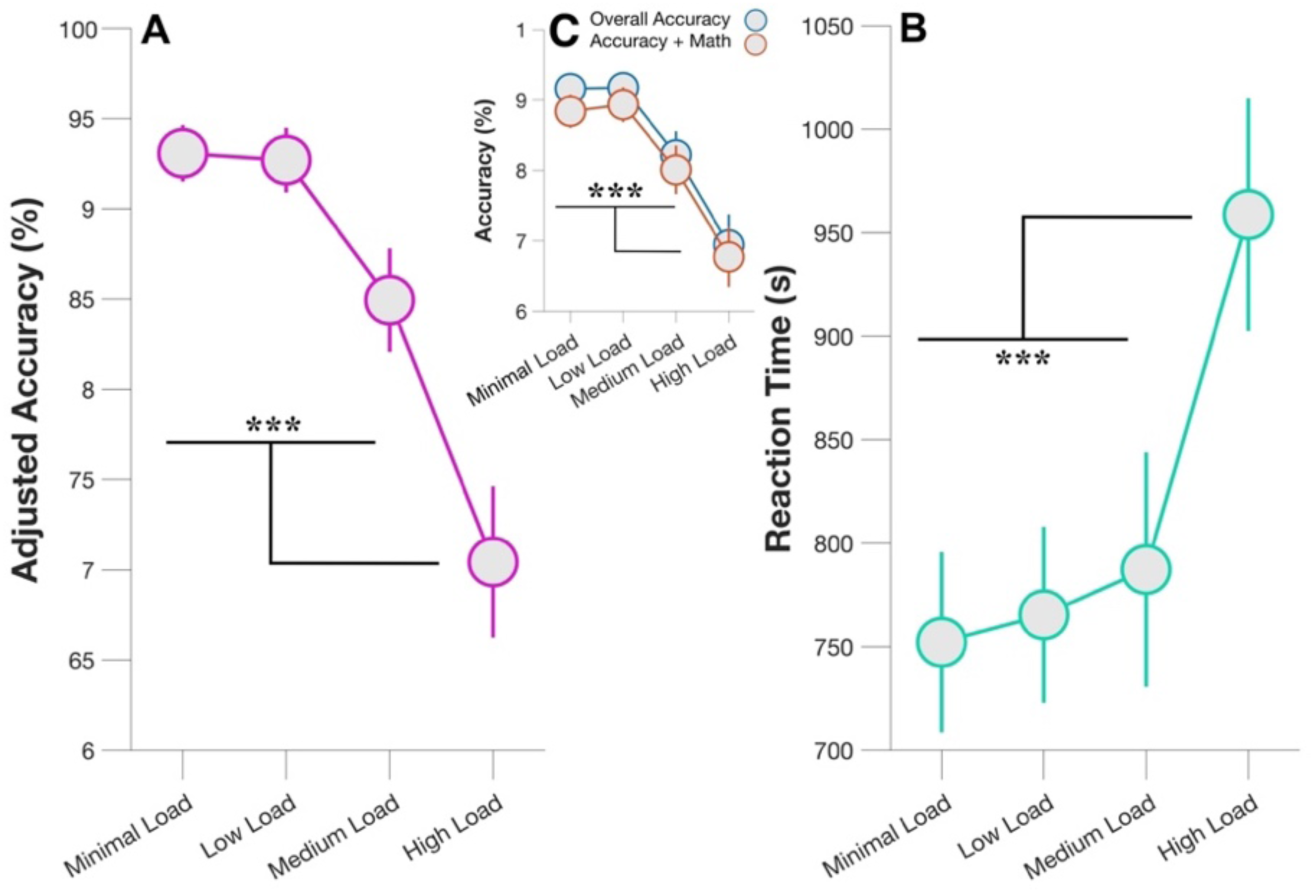
Behavioural Performance on the DMT after outlier removal. Panel A illustrates task accuracy, while panel B depicts reaction time (RT). In panel A, the y-axis represents adjusted accuracy (%), which accounts for trials where participants correctly solved the DMT and arithmetic task, while the x-axis displays the four cognitive load conditions (difficulty levels). Panel C provides an overview of overall accuracy, including both “all responses” and “accuracy + math,” the latter specifically considering trials where participants correctly responded to arithmetic task. The error bars represent ±1 SEM.

### Experiment 2 (Main)

Wilcoxon signed-rank tests revealed consistent effects of load level across conditions. As shown in Figure 2, Panel A and B, participants were significantly slower in the HL compared to LL condition, (no nudge; Z = –6.429, *p* < 0.001), and accuracy was also significantly reduced, (Z = 7.532, *p* < 0.001). Similarly, when comparing within-subject conditions for nudge (D), significant differences emerged for both RT (Z = –4.793, *p* < 0.001) and accuracy (Z = 4.323, *p* < 0.001). Finally, for nudge (A), significant effects of load were again observed for RT (Z = –4.533, *p* < 0.001) and accuracy (Z = 5.089, *p* < 0.001).

A two-way mixed ANOVA was conducted to examine the effects of nudge (between-subject factor: no nudge, nudge (D), and nudge (A)) and load (within-subject factor: NL, LL, and HL) on audit task RTs. A significant main effect of nudge on RTs was observed, (F (2, 111) = 225.20, *p* < 0.001, η² = 0.80). However, the main effect of load was not significant, (F (2, 222) = 1.39, *p* = 0.25, η² = 0.012). The nudge × load interaction was significant, (F (4, 222) = 24.75, *p* < 0.001, η² = 0.31) (Figure 2, Panel C).

Similarly, for accuracy, while the main effect of nudge was significant, (F (2, 111) = 98.33, *p* < 0.001, η² = 0.64), the main effect of load was not significant, (F (2, 222) = 0.15, *p* = 0.86, η² = 0.001). The nudge × load interaction was significant, (F (4, 222) = 17.52, *p* < 0.001, η² = 0.24) (Figure 2, Panel D).

**Figure 2.**
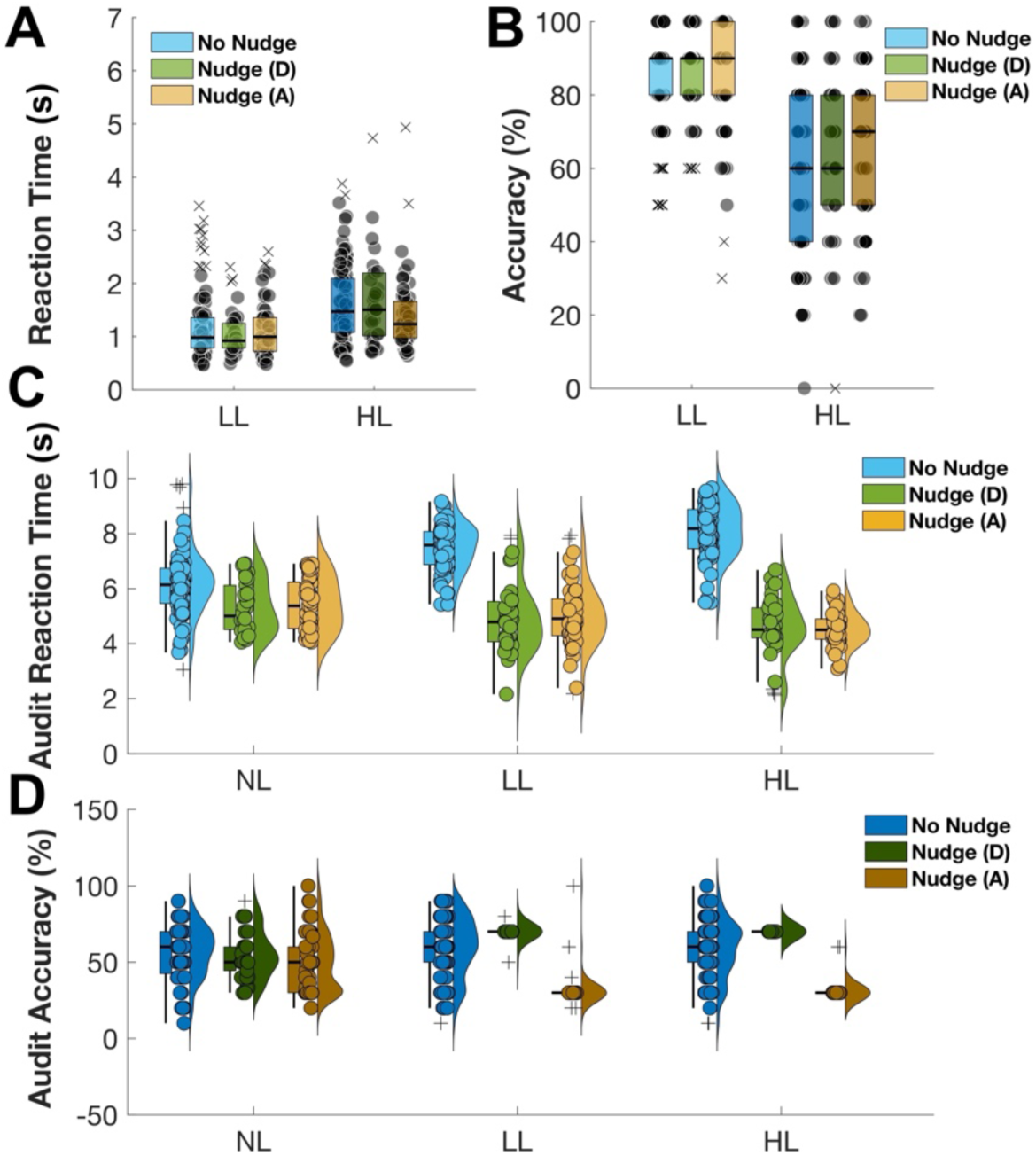
Panels A and B depict the behavioural performance for the DMT, showing RT (ms) and accuracy rate (%), respectively. Panels C and D illustrate RT (ms) and accuracy rate (%) for the audit task. In panels A and B, each whisker represents the interquartile range (IQR), the horizontal black line denotes the mean, and black dots represent individual data points. In panels C and D, whiskers indicate the IQR, horizontal black lines show the mean, and error bars are included. Circles represent individual data points, while the kernel density plots visualize the data distribution. distribution.

The pattern of results remained consistent with those obtained from the full dataset, indicating that the observed effects were not driven by extreme values.

Analysis on accuracy + math and overall accuracy as well as pairwise comparisons with adjusted *p*-values were not repeated in the supplementary analyses, as these were already reported in the main manuscript. The supplementary section is intended to provide robustness checks, not to duplicate corrected inferential results.

## References

Anderson, S. F., Kelley, K., & Maxwell, S. E. (2017). Sample-size planning for more accurate statistical power: A method adjusting sample effect sizes for publication bias and uncertainty. Psychological science, 28(11), 1547–1562. 10.1177/095679761772372

Anwyl-Irvine, A. L., Massonnie, J., Flitton, A., Kirkham, N., & Evershed, J. K. (2020). Gorilla in our midst: An online behavioral experiment builder. Behav Res Methods, 52(1), 388–407. 10.3758/s13428-019-01237-x

Arkes, H. R., Dawes, R. M., & Christensen, C. (1986). Factors influencing the use of a decision rule in a probabilistic task. Organizational behavior and human decision processes, 37(1), 93–110.

Asare, S. K., Trompeter, G. M., & Wright, A. M. (2000). The effect of accountability and time budgets on auditors’ testing strategies. Contemporary Accounting Research, 17(4), 539–560. 10.1506/F1EG-9EJG-DJ0B-JD32

Austin, A. A. (2023). Remembering fraud in the future: Investigating and improving auditors’ attention to fraud during audit testing. Contemporary Accounting Research, 40(2), 925–951. 10.1111/1911-3846.12843

Backs, R. W., & Seljos, K. A. (1994). Metabolic and cardiorespiratory measures of mental effort: the effects of level of difficulty in a working memory task. International Journal of psychophysiology, 16(1), 57–68. 10.1016/0167-8760(94)90042-6

Bago, B., & De Neys, W. (2017). Fast logic?: Examining the time course assumption of dual process theory. Cognition, 158, 90–109. 10.1016/j.cognition.2016.10.014

Bago, B., Rand, D. G., & Pennycook, G. (2020). Fake news, fast and slow: Deliberation reduces belief in false (but not true) news headlines. Journal of experimental psychology: General, 149(8), 1608. 10.1037/xge0000729

Barrouillet, P., Bernardin, S., & Camos, V. (2004). Time constraints and resource sharing in adults’ working memory spans. Journal of experimental psychology: General, 133(1), 83. 10.1037/0096-3445.133.1.83

Barrouillet, P., Bernardin, S., Portrat, S., Vergauwe, E., & Camos, V. (2007). Time and cognitive load in working memory. J Exp Psychol Learn Mem Cogn, 33(3), 570–585. 10.1037/0278-7393.33.3.570

Barrouillet, P., & Camos, V. (2012). As time goes by: Temporal constraints in working memory. Current directions in psychological science, 21(6), 413–419. 10.1177/0963721412459513

Barzykowski, K., & Niedzwienska, A. (2018). Involuntary autobiographical memories are relatively more often reported during high cognitive load tasks. Acta Psychol (Amst), 182, 119–128. 10.1016/j.actpsy.2017.11.014

Baumeister, R. F. (2001). Ego depletion, the executive function, and self-control: An energy model of the self in personality. 10.1080/152988602317319302

Bedeir, R. E. (2024). The differential impact of distracted auditors in managing portfolio of financially distressed audit clients on audit quality: the role of professional skepticism. Future Business Journal, 10(1), 36. 10.1186/s43093-024-00321-9

Bethell-Fox, C. E., & Shepard, R. N. (1988). Mental rotation: Effects of stimulus complexity and familiarity. Journal of Experimental Psychology: Human Perception and Performance, 14(1), 12. 10.1037/0096-1523.14.1.12

Black, V., Allen, J., Aazh, H., Johnson, S. L., & Erfanian, M. (2025). Misophonia Symptoms Severity is Attributed to Impaired Flexibility and Heightened Rumination. BioRxiv, 2025.2001.2013.632542. 10.1101/2025.01.13.632542

Board, P. C. A. O. (2012). Maintaining and Applying Professional Skepticism in Audits. Staff Audit Practice Alert No. 10. In: PCAOB Washington, DC.

Bohn, R., & Short, J. E. (2012). Info capacity| measuring consumer information. International Journal of Communication, 6, 21.

Botvinick, M. M., Cohen, J. D., & Carter, C. S. (2004). Conflict monitoring and anterior cingulate cortex: an update. Trends Cogn Sci, 8(12), 539–546. 10.1016/j.tics.2004.10.003

Brunye, T. T., Martis, S. B., & Taylor, H. A. (2018). Cognitive load during route selection increases reliance on spatial heuristics. Q J Exp Psychol (Hove), 71(5), 1045–1056. 10.1080/17470218.2017.1310268

Bulus, M. (2023). Pwrss: Statistical power and sample size calculation tools. R package version 0.3, 1.

Camos, V., & Portrat, S. (2015). The impact of cognitive load on delayed recall. Psychon Bull Rev, 22(4), 1029–1034. 10.3758/s13423-014-0772-5

Cohen, J. (1988). Statistical power analysts for the behavioral sciences. 2nd edn Hillsdale. NJL Erlbaum Associates.

Correll, J., Mellinger, C., McClelland, G. H., & Judd, C. M. (2020). Avoid Cohen’s ‘small’,‘medium’, and ‘large’for power analysis. Trends in cognitive sciences, 24(3), 200–207. 10.1016/j.tics.2019.12.009

Croskerry, P. (2003). The importance of cognitive errors in diagnosis and strategies to minimize them. Academic medicine, 78(8), 775–780. 10.1097/00001888-200308000-00003

Csipo, T., Lipecz, A., Mukli, P., Bahadli, D., Abdulhussein, O., Owens, C. D., Tarantini, S., Hand, R. A., Yabluchanska, V., & Kellawan, J. M. (2021). Increased cognitive workload evokes greater neurovascular coupling responses in healthy young adults. PLoS One, 16(5), e0250043. 10.1371/journal.pone.0250043

Evans, J. S., & Stanovich, K. E. (2013). Dual-Process Theories of Higher Cognition: Advancing the Debate. Perspect Psychol Sci, 8(3), 223–241. 10.1177/1745691612460685

Evans, J. S. B., & Curtis-Holmes, J. (2005). Rapid responding increases belief bias: Evidence for the dual-process theory of reasoning. Thinking & Reasoning, 11(4), 382–389. 10.1080/13546780542000005

Gailliot, M. T. (2008). Unlocking the Energy Dynamics of Executive Functioning: Linking Executive Functioning to Brain Glycogen. Perspect Psychol Sci, 3(4), 245–263. 10.1111/j.1745-6924.2008.00077.x

Gajewski, J.-F., Heimann, M., Léger, P.-M., & Teye, P. (2024a). Enhancing auditors’ professional skepticism through nudges: an eye-tracking experiment. Accounting and Business Research, 1–19. 10.1080/00014788.2024.2364215

Gajewski, J.-F., Heimann, M., & Meunier, L. (2022). Nudges in SRI: the power of the default option. Journal of Business Ethics, 1–20. 10.1007/s10551-020-04731-x

Gajewski, J.-F., Heimann, M., Meunier, L., & Ohadi, S. (2024b). Nudges for responsible finance? A survey of interventions targeted at financial decision making. Corporate Social Responsibility and Environmental Management, 31(2), 1203–1219. 10.1002/csr.2625

Gigerenzer, G. (2008). Why Heuristics Work. Perspect Psychol Sci, 3(1), 20–29. 10.1111/j.1745-6916.2008.00058.x

Gigerenzer, G., & Edwards, A. (2003). Simple tools for understanding risks: from innumeracy to insight. Bmj, 327(7417), 741–744. 10.1136/bmj.327.7417.741

Gigerenzer, G., & Gaissmaier, W. (2011). Heuristic decision making. Annual review of psychology, 62(1), 451–482. 10.1146/annurev-psych-120709-145346

Hauser, D. J., & Schwarz, N. (2016). Attentive Turkers: MTurk participants perform better on online attention checks than do subject pool participants. Behavior Research Methods, 48(1), 400–407. 10.3758/s13428-015-0578-z

Heyes, C. (2018). Cognitive gadgets: The cultural evolution of thinking. Harvard University Press.

Hu, J., Xin, C., Zhang, M., & Chen, Y. (2024). The effect of cognitive load and time stress on prospective memory and its components. Current Psychology, 43(2), 1670–1684. 10.1007/s12144-023-04354-1

Hurtt, R. K. (2010). Development of a scale to measure professional skepticism. Auditing: A Journal of Practice & Theory, 29(1), 149–171. 10.2308/aud.2010.29.1.149

Hurtt, R. K., Brown-Liburd, H., Earley, C. E., & Krishnamoorthy, G. (2013). Research on auditor professional skepticism: Literature synthesis and opportunities for future research. Auditing: A Journal of Practice & Theory, 32(Supplement 1), 45–97. 10.2308/ajpt-50361

Jachimowicz, J. M., Duncan, S., Weber, E. U., & Johnson, E. J. (2019). When and why defaults influence decisions: A meta-analysis of default effects. Behavioural Public Policy, 3(2), 159–186. 10.1017/bpp.2018.43

Jeffreys, H. (1961). The Theory of Probability. 1998 ed. In: OUP Oxford. Google-Books-ID: vh9Act9rtzQC.

Johari, R. J., Ridzoan, N. S., & Zarefar, A. (2019). The influence of work overload, time pressure and social influence pressure on auditors’ job performance. International Journal of Financial Research, 10(3), 88–106. 10.5430/ijfr.v10n3p88

Kahneman, D., & Klein, G. (2009). Conditions for intuitive expertise: a failure to disagree. American psychologist, 64(6), 515. 10.1037/a0016755

Kahneman, D., Sibony, O., & Sunstein, C. R. (2021). Noise: A flaw in human judgment. Hachette UK.

Kool, W., & Botvinick, M. (2018). Mental labour. Nature human behaviour, 2(12), 899–908. 10.1038/s41562-018-0401-9

Kurzban, R., Duckworth, A., Kable, J. W., & Myers, J. (2013). An opportunity cost model of subjective effort and task performance. Behavioral and Brain Sciences, 36(6), 661–679. 10.1017/S0140525X12003196

Lakens, D. (2022). Sample size justification. Collabra: psychology, 8(1), 33267.

Lambert, T. A., Jones, K. L., Brazel, J. F., & Showalter, D. S. (2017). Audit time pressure and earnings quality: An examination of accelerated filings. Accounting, Organizations and Society, 58, 50–66. 10.1016/j.aos.2017.03.003

Leung, H. C., Seelig, D., & Gore, J. C. (2004). The effect of memory load on cortical activity in the spatial working memory circuit. Cogn Affect Behav Neurosci, 4(4), 553–563. 10.3758/cabn.4.4.553

Lewandowsky, S., Ecker, U. K., & Cook, J. (2017). Beyond misinformation: Understanding and coping with the “post-truth” era. Journal of applied research in memory and cognition, 6(4), 353–369. 10.1016/j.jarmac.2017.07.008

Li, H. L., & van Rossum, M. C. (2020). Energy efficient synaptic plasticity. Elife, 9. 10.7554/eLife.50804

Mansouri, Koechlin, E., Rosa, M. G. P., & Buckley, M. J. (2017). Managing competing goals - a key role for the frontopolar cortex. Nat Rev Neurosci, 18(11), 645–657. 10.1038/nrn.2017.111

Marchiori, D. R., Adriaanse, M. A., & De Ridder, D. T. (2017). Unresolved questions in nudging research: Putting the psychology back in nudging. Social and personality psychology compass, 11(1), e12297. 10.1111/spc3.12297

MATLAB. (2019). Release, Statistics Toolbox, The MathWorks, Inc., Natick, Massachusetts, United States. In: ed.

Mazzoni, G. (2019). Involuntary memories and involuntary future thinking differently tax cognitive resources. Psychol Res, 83(4), 684–697. 10.1007/s00426-018-1123-3

McDaniel, L. S. (1990). The effects of time pressure and audit program structure on audit performance. Journal of accounting research, 28(2), 267–285. 10.2307/2491150

Mercier, H., & Sperber, D. (2011). Why do humans reason? Arguments for an argumentative theory. Behavioral and Brain Sciences, 34(2), 57–74. 10.1017/S0140525X10000968

Mercier, H., & Sperber, D. (2017). The enigma of reason. Harvard University Press.

Mertens, S., Herberz, M., Hahnel, U. J., & Brosch, T. (2022). The effectiveness of nudging: A meta-analysis of choice architecture interventions across behavioral domains. Proc. Natl. Acad. Sci. 10.1073/pnas.2107346118

Meunier, L., Bashirzadeh, Y., & Ohadi, S. (2024a). Framing the Default Option Right. Journal of Behavioral Decision Making, 37(3), e2395. 10.1002/bdm.2395

Meunier, L., & Richit, S. (2024b). Testing four nudges in socially responsible investments: Default winner by inertia. Business ethics, the environment & responsibility, 33(3), 392–415. 10.1111/beer.12612

Miller, J. (1991). Reaction time analysis with outlier exclusion: bias varies with sample size. Q J Exp Psychol A, 43(4), 907–912. 10.1080/14640749108400962

Miller, J. (2023). Outlier exclusion procedures for reaction time analysis: The cures are generally worse than the disease. J Exp Psychol Gen, 152(11), 3189–3217. 10.1037/xge0001450

Miyake, A., Friedman, N. P., Rettinger, D. A., Shah, P., & Hegarty, M. (2001). How are visuospatial working memory, executive functioning, and spatial abilities related? A latent-variable analysis. J Exp Psychol Gen, 130(4), 621–640. 10.1037//0096-3445.130.4.621

Morey, R. D., & Rouder, J. (2015). Using the BayesFactor package (version 0.9. 2). In.

Münscher, R., Vetter, M., & Scheuerle, T. (2016). A review and taxonomy of choice architecture techniques. Journal of Behavioral Decision Making, 29(5), 511–524. 10.1002/bdm.1897

Nelson, M. W. (2009). A model and literature review of professional skepticism in auditing. Auditing: A Journal of Practice & Theory, 28(2), 1–34. 10.2308/aud.2009.28.2.1

Ninomiya, Y., Iwata, T., Terai, H., & Miwa, K. (2024). Effect of cognitive load and working memory capacity on the efficiency of discovering better alternatives: A survival analysis. Mem Cognit, 52(1), 115–131. 10.3758/s13421-023-01448-w

Normand, M. P. (2008). Science, skepticism, and applied behavior analysis. Behav Anal Pract, 1(2), 42–49. 10.1007/BF03391727

Ordali, E., Marcos-Prieto, P., Avvenuti, G., Ricciardi, E., Boncinelli, L., Pietrini, P., Bernardi, G., & Bilancini, E. (2024). Prolonged exertion of self-control causes increased sleep-like frontal brain activity and changes in aggressivity and punishment. Proc Natl Acad Sci U S A, 121(47), e2404213121. 10.1073/pnas.2404213121

Otley, D. T., & Pierce, B. J. (1996). The operation of control systems in large audit firms. Auditing, 15(2), 65.

Padamsey, Z., & Rochefort, N. L. (2023). Paying the brain’s energy bill. Curr Opin Neurobiol, 78, 102668. 10.1016/j.conb.2022.102668

Payne, J. W., Bettman, J. R., & Johnson, E. J. (1993). The adaptive decision maker. Cambridge university press.

PCAOB. (2022). Staff Update and Preview of 2021 Inspection Observations. https://pcaob-assets.azureedge.net/pcaob-dev/docs/default-source/documents/staff-preview-2021-inspection-observations-spotlight.pdf?sfvrsn=d2590627_2/

PCAOB. (2023). Professional Competence and Skepticism are Essential to Quality Audits. Retrieved from Available from: https://pcaobus.org/documents/competence-and-skepticism-spotlight.pdf(open in a new window)

Peer, E., Brandimarte, L., Samat, S., & Acquisti, A. (2017). Beyond the Turk: Alternative platforms for crowdsourcing behavioral research. Journal of experimental social psychology, 70, 153–163. 10.1016/j.jesp.2017.01.006

Pennycook, G., & Rand, D. G. (2019). Lazy, not biased: Susceptibility to partisan fake news is better explained by lack of reasoning than by motivated reasoning. Cognition, 188, 39–50. 10.1016/j.cognition.2018.06.011

Phillips, F. (1999). Auditor attention to and judgments of aggressive financial reporting. Journal of accounting research, 37(1), 167–189. 10.2307/2491402

Raichle, M. E., & Gusnard, D. A. (2002). Appraising the brain’s energy budget. Proc Natl Acad Sci U S A, 99(16), 10237–10239. 10.1073/pnas.172399499

Reijula, S., & Hertwig, R. (2022). Self-nudging and the citizen choice architect. Behavioural Public Policy, 6(1), 119–149. 10.1017/bpp.2020.5

Rose, J. M. (2007). Attention to evidence of aggressive financial reporting and intentional misstatement judgments: Effects of experience and trust. Behavioral Research in Accounting, 19(1), 215–229. 10.2308/bria.2007.19.1.215

Rouder, J. N., Morey, R. D., Speckman, P. L., & Province, J. M. (2012). Default Bayes factors for ANOVA designs. Journal of mathematical psychology, 56(5), 356–374. 10.1016/j.jmp.2012.08.001

Rouder, J. N., Speckman, P. L., Sun, D., Morey, R. D., & Iverson, G. (2009). Bayesian t tests for accepting and rejecting the null hypothesis. Psychonomic bulletin & review, 16, 225–237. 10.3758/PBR.16.2.225

Salthouse, T. A. (2009). When does age-related cognitive decline begin? Neurobiology of aging, 30(4), 507–514. 10.1016/j.neurobiolaging.2008.09.023

Salthouse, T. A., Pink, J. E., & Tucker-Drob, E. M. (2008). Contextual analysis of fluid intelligence. Intelligence, 36(5), 464–486. 10.1016/j.intell.2007.10.003

Shanteau, J. (1992). Competence in experts: The role of task characteristics. Organizational behavior and human decision processes, 53(2), 252–266. 10.1016/0749-5978(92)90064-E

Simon, J. R., & Wolf, J. D. (1963). Choice reaction time as a function of angular stimulus-response correspondence and age. Ergonomics, 6(1), 99–105. 10.1080/00140136308930679

Sisakhti, M., Sachdev, P. S., & Batouli, S. A. H. (2021). The effect of cognitive load on the retrieval of long-term memory: An fMRI study. Frontiers in human neuroscience, 15, 700146. 10.3389/fnhum.2021.700146

Sliwinski, M. J., Mogle, J. A., Hyun, J., Munoz, E., Smyth, J. M., & Lipton, R. B. (2018). Reliability and validity of ambulatory cognitive assessments. Assessment, 25(1), 14–30. 10.1177/1073191116643164

Sperber, D., Clément, F., Heintz, C., Mascaro, O., Mercier, H., Origgi, G., & Wilson, D. (2010). Epistemic vigilance. Mind & language, 25(4), 359–393. 10.1111/j.1468-0017.2010.01394.x

Stanovich, K. (2011). Rationality and the reflective mind. Oxford University Press. 10.1093/acprof:oso/9780195341140.001.0001

Stanovich, K. E., West, R. F., & Toplak, M. E. (2016). The rationality quotient: Toward a test of rational thinking. MIT press.

Steffel, M., Williams, E. F., & Pogacar, R. (2016). Ethically deployed defaults: Transparency and consumer protection through disclosure and preference articulation. Journal of Marketing Research, 53(5), 865–880.

Sterling, P., & Laughlin, S. (2015). Principles of neural design. MIT press.

Sunstein, C. R. (2023). Eight misconceptions about nudges. In Research Handbook on Nudges and Society (pp. 319–328). Edward Elgar Publishing.

Svanström, T. (2016). Time pressure, training activities and dysfunctional auditor behaviour: evidence from small audit firms. International Journal of Auditing, 20(1), 42–51. 10.1111/ijau.12054

Sweller, J. (1988). Cognitive load during problem solving: Effects on learning. Cognitive science, 12(2), 257–285. 10.1207/s15516709cog1202_4

Team, R. C. (2023). R: A Language and Environment for Statistical Computing. R Foundation for Statistical Computing, Vienna. https://www.R-project.org/

Thaler, R. H., & Sunstein, C. R. (2008). Nudge: Improving Decisions About Health. Wealth, and Happiness, 3.

Van Gestel, L., Adriaanse, M., & De Ridder, D. (2021). Do nudges make use of automatic processing? Unraveling the effects of a default nudge under type 1 and type 2 processing. Comprehensive Results in Social Psychology, 5(1-3), 4–24. 10.1080/23743603.2020.1808456

Vannucci, M., Pelagatti, C., Hanczakowski, M., Mazzoni, G., & Paccani, C. R. (2015). Why are we not flooded by involuntary autobiographical memories? Few cues are more effective than many. Psychol Res, 79(6), 1077–1085. 10.1007/s00426-014-0632-y

Vrij, A., Fisher, R. P., & Blank, H. (2017). A cognitive approach to lie detection: A meta-analysis. Legal and Criminological Psychology, 22(1), 1–21. 10.1111/lcrp.12088

Westbrook, A., & Braver, T. S. (2015). Cognitive effort: A neuroeconomic approach. Cognitive, Affective, & Behavioral Neuroscience, 15, 395–415. 10.3758/s13415-015-0334-y

Wiehler, A., Branzoli, F., Adanyeguh, I., Mochel, F., & Pessiglione, M. (2022). A neuro-metabolic account of why daylong cognitive work alters the control of economic decisions. Curr Biol, 32(16), 3564–3575 e3565. 10.1016/j.cub.2022.07.010

Zhang, B., Zhang, J. X., Huang, S., Kong, L., & Wang, S. (2011). Effects of load on the guidance of visual attention from working memory. Vision Res, 51(23-24), 2356–2361. 10.1016/j.visres.2011.09.008

Zhao, X., Chen, A., & West, R. (2010). The influence of working memory load on the Simon effect. Psychon Bull Rev, 17(5), 687–692. 10.3758/PBR.17.5.687

